# Neural Activation in the Olfactory Epithelium of the East African Cichlid (*Haplochromis chilotes*) in Response to Odorant Exposure

**DOI:** 10.1101/2021.05.25.445711

**Authors:** Riki Kawamura, Masato Nikaido

**Affiliations:** School of Life Science and Technology, Tokyo Institute of Technology

**Keywords:** *c-fos*, V1R, V2R, amino acid, urine collection

## Abstract

Fishes use olfaction to gain various information vital for survival and communication. To understand biodiversity in fishes, it is important to identify what receptors individual fish use to detect specific chemical compounds. However, studies of fish olfactory receptors and their ligands are still limited to a few model organisms represented primarily by zebrafish. Here, we tested the neural activation of olfactory sensory neurons (OSNs) in East African cichlids, the most diversified teleost lineage, by *in situ* hybridization with a *c-fos* riboprobe. We confirmed that microvillous neurons contributed the most to the detection of amino acids, as in other fishes. Conversely, we found that ciliated neurons contributed the most to detection of conjugated steroids, known as pheromone candidates. We also found that V2Rs, the major expressed receptor in microvillous neurons exhibited differential responsiveness to amino acids, and further suggested that the cichlid-specific duplication of *V2R* led to ligand differentiation in cichlids by demonstrating a differential response to arginine. Finally, we established a nonlethal method to collect cichlid urine and showed how various OSNs, including *V1R*^+^ neurons, respond to male urine. This study provides an experimental basis for understanding how cichlids encode natural odors and illuminates how olfaction has contributed to the diversification of cichlids by combining with phylogenetic studies of olfactory receptors gene evolutions.

## Introduction

Animals use olfaction to guide many complex behaviors for improving fitness and survival, such as foraging for food, choosing mates, recognizing territories, migrating, and avoiding predators. Olfactory receptors play a vital role in these behaviors. Hence, to understand the specific ecology of these genes, their receptor function must be revealed.

In fishes, several soluble chemical compounds are detected by the olfactory epithelium (OE) as odor. Several odorants are known to drive feeding behavior in fishes. For example, amino acids drives feeding behavior in zebrafish and salmonid species (Valentinčič *et al*., 1999; Hara, 2006; Koide *et al*., 2009), polyamines in goldfish (Rolen *et al*., 2003), and nucleotides in zebrafish (Wakisaka et al., 2017). These odorants sometimes work differently in other species, e.g. amino acids works as migration pheromones or sex pheromones in salmonid species (Shoji *et al*., 2003; Yambe *et al*., 2006; Yamamoto *et al*., 2010, 2013), and polyamines drive aversive behavior in zebrafish (Hussain et al., 2013). Sex steroids and prostaglandins have been shown to act as pheromones in goldfish (Dulka et al., 1987; Sorensen et al., 1988, 2005; Stacey et al., 1989) and prostaglandins in zebrafish (Yabuki et al., 2016). Bile acids are detected by several species, and they drive migration behavior in lamprey, although their functions in other fishes remain controversial (Li *et al*., 1995; Michel and Lubomudrov, 1995; Zhang *et al*., 2001; Huertas *et al*., 2010).

Fishes detect these odorants by olfactory receptors, which are encoded by four G protein-coupled receptor multigene families: odorant receptor (OR; Buck and Axel, 1991), trace amine-associated receptor (TAAR; Liberles and Buck, 2006), vomeronasal type-1 receptor (V1R, also termed as ORA; Dulac and Axel, 1995), and vomeronasal type-2 receptor (V2R, also termed as OlfC; Herrada and Dulac, 1997). Single GPCR A2c are also found in the zebrafish OE (Wakisaka et al., 2017). There are also some putatively expressed olfactory receptor families such as formyl peptide receptors (FPRs, Rivière et al., 2009) and membrane-spanning 4A receptors (MS4As, Greer et al., 2016) which are reported to be expressed in mammal olfactory-related organs.

The above receptors are expressed in several types of olfactory sensory neurons (OSNs). Ciliated neurons (Hansen *et al*., 2004; Sorensen and Sato, 2005), the major OSNs expressing OR or TAAR, detect a broad range of odorants such as amino acids, sex steroids, prostaglandins, and bile acids (Sato and Suzuki, 2001; Hansen *et al*., 2003; Sato and Sorensen, 2018). Furthermore, some studies on zebrafish redefined OR114 receptor as the fish pheromone prostaglandin F_2α_ receptor (Yabuki *et al*., 2016), and also several TAAR receptors as polyamine receptors (Hussain et al., 2013; Li et al., 2015). On the other hand, microvillous neurons, another major group of OSN expresses V1R or V2R, specifically detect amino acids (Sato and Suzuki, 2001; Hansen *et al*., 2003; Sato and Sorensen, 2018). Two studies also suggest that V2R receptors detect amino acids (Koide *et al*., 2009; DeMaria *et al*., 2013). Moreover, several heterologous expression experiments show that zebrafish V1R receptors bind 4-hydroxyphenyl acetate (4HPAA) and bile acids (Behrens *et al*., 2014; Cong *et al*., 2019). Pear-shaped neurons, which express A2C, detect adenosine (Wakisaka et al., 2017). It is not yet known what is detected by other minor OSNs such as kappe neurons (Ahuja et al., 2015), and crypt neurons (Oka *et al*., 2012), which express single V1R (V1R4/ora4).

In general, relatively few olfactory receptors have been studied and these studies are limited to the zebrafish model and only several nonmodel organisms such as salmon and goldfish. Alternatively, the clade Neoteleostei, which includes approximately 60% of fish taxonomic diversity, has not been the focus for studies of olfaction. Hence, investigating this large diverse clade of nonmodel species remains crucial for understanding fish evolution.

In this study, we focused on cichlids, one of the most diversified lineages of vertebrates. Cichlids in the East African Great Lakes represent one of the most striking examples of vertebrate adaptive radiation (Kocher, 2004). Because of the highly diversified nuptial colouration, the visual ecology of cichlids has been important area of research demonstrating the importance of vision in driving cichlid speciation (Terai et al., 2006; Seehausen et al., 2008). Although olfaction has drawn less attention from cichlid biologists compared to vision, cichlids also utilize olfaction in many different ecological contexts (Keller-Costa *et al*., 2015). For example, olfaction contributes to conspecific recognition of *Pseudotropheus emmiltos* (Plenderleith *et al*., 2005) and sexual imprinting of *Pundamilia* species (Verzijden and Ten Cate, 2007). Other studies show that male tilapia evaluate the sexual status of potential mates from female urine (Miranda et al., 2005), and that a glucuronidated steroid in male tilapia urine works as a priming pheromone (Keller-Costa et al., 2014). Moreover, we previously found several highly diverse polymorphic alleles in the V1R receptors of East African cichlids (Nikaido *et al*., 2014), and the copy number of V2R receptors has increased in East African cichlid genomes (Nikaido *et al*., 2013), which suggests the functional importance of olfaction potentially driving cichlid adaptive radiation. However, like other fishes, little is understood about the ligand specificity of individual OSNs in cichlids.

Here, we performed *in situ* hybridization with a riboprobe of neural activity marker gene *c-fos* to investigate OE in the East African cichlid *Haplochromis chilotes*. We tested the response of several types of OSNs, i.e. microvillous neurons, *V2Rs*^+^ neurons and *V1Rs*^+^ neurons, and reported the ligand specificity of several odorants. We also tested several individual V2Rs for several amino acids and V1Rs for cichlid male urine. This study offers important initial experimental insights into our fundamental understanding of how cichlids encode natural odors. Moreover, it reveals how olfaction has contributed to the diversification of cichlids through phylogenetic studies of olfactory receptor gene evolution.

## Materials and Methods

### Fish

Cichlids (*Haplochromis chilotes*) were maintained at 27°C on a 12 h light/12 h dark cycle. Six to twelve individuals were kept in a plastic tank (40 cm × 25 cm × 36 cm). Mature males were used for experiments.

### Odorant solutions

Twenty proteinogenic amino acids (arginine, histidine, lysine, aspartate, glutamate, serine, threonine, asparagine, glutamine, cysteine, glycine, proline, alanine, isoleucine, leucine, methionine, phenylalanine, tryptophan, tyrosine, and valine), 4HPAA, and lithocholic acid (LCA) were purchased from Wako Pure Chemical Industries and Sigma Chemical Co. Each amino acid (except tyrosine) and 4HPAA was dissolved in ultrapure water to 12 mM to create a stock solutions. Tyrosine and LCA were dissolved in 6 mM NaOH aqueous solution to 12 mM as the stock solutions. Three conjugated steroids, dehydroepiandrosterone 3-sulfate (DHEA-s), β-estradiol 17-(β-D-glucuronide) (E_2_-17g), and β-estradiol 3,17-disulfate (E_2_-3, 17s) were respectively purchased from Tokyo Chemical Industry, Cayman Chemical Co., and Santa Cruz Biotechnology, respectively, and dissolved in DMSO to 10 mM to create a stock solutions. Food extract was prepared by the following procedure: First, 2 g of crushed fish food Otohime EP1 (Marubeni Nisshin Feed Co.) was added to ultrapure water up to 14 mL and vortexed. After incubating at room temperature for 5 min, the extraction liquid was centrifuged at 8,000 × g for 5 min, and the supernatant was collected as the food extract stock solution. Each stock solution was stored at 4°C until the exposure experiment. Stock solutions was diluted with ultrapure water prior to the exposure, and 15 mL diluted solution was applied experimentally. Each solution was diluted as follows: the mixture of 20 amino acids/amino acids group A-D was diluted to 400 µM (final concentration at 2 µM in the exposure tank); the mixture of conjugated steroids was diluted to 6.6 µM each (final concentration at 33 nM in the exposure tank); arginine/lysin/glutamate/aspartate/4HPAA was diluted to 2 mM (final concentration at 10 µM in the exposure tank); LCA was diluted to 4 mM (final concentration at 20 µM in the exposure tank); food extract was diluted 75-fold (final concentration at 15,000-fold dilution in the exposure tank).

### Urine collection

Urine was collected from mature male cichlids. Although several studies have collected nondiluted urine from fish, they were limited to larger species such as Masu salmon, rainbow trout, Mozambique tilapia, and Senegalese sole (Yambe *et al*., 1999; Sato and Suzuki, 2001; Keller-Costa *et al*., 2014; Fatsini *et al*., 2017). By adapting methods to collect urine from Masu salmon employed in Yambe *et al*., 1999, we developed a nonlethal methods to collect urine directly from cichlids, whose size is approximately 6–9 cm under swimming conditions (Figure 1A, B). We used a dental root canal cleaning probe needle (28G, 490703, BSA Sakurai Co.) to construct a sampling catheter (Figure 1A). This needle has a hole in the side near the tip to prevent clogging. Approximately 0.5–1.2 cm from the tip, the needle was gently bent approximately 90° so that the hollow tube structure remained open (Figure 1A). This bent needle was connected to a 15 mL centrifuge tube using silicon tubing (OD: 10 mm, ID: 0.5 mm) fixed with adhesive (Aron Alpha EXTRA Fast-Acting Versatile, Konishi) to trap the urine. The centrifuge tube was then further connected to an aspirator (DAS-01, As one) to aspirate the urine.

**Figure 1.**
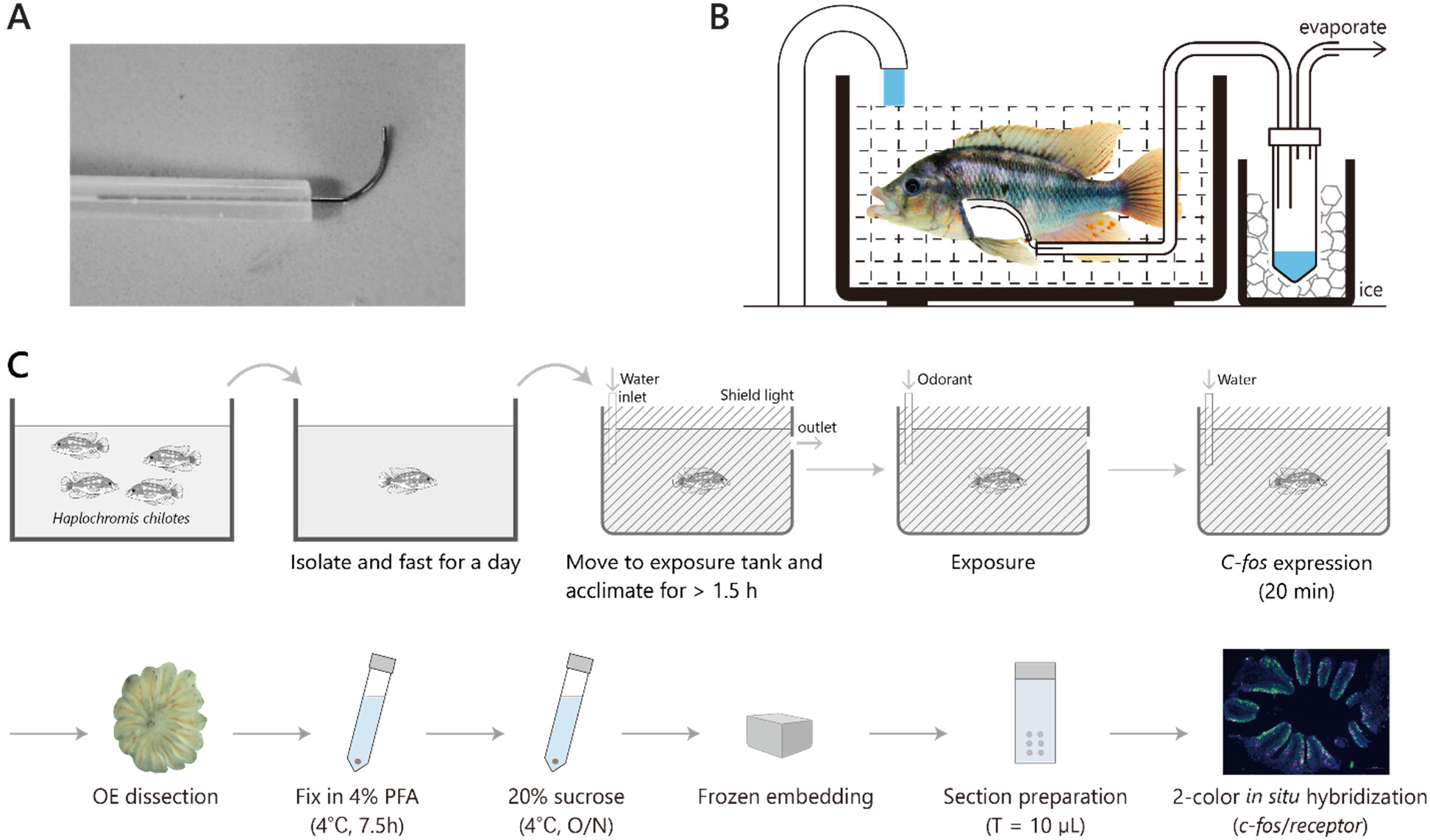
Schematic drawing of methods. **(A)** Catheter for urine collection. The catheter has a hole in the side near the tip of the needle to prevent clogging. **(B)** Schematic drawing of nonlethal urine collection protocol. **(C)** Schematic workflow for detection of odorant-induced neural response. Cichlids were exposed to an odorant. Olfactory epithelia were then isolated, fixed, and frozen embedded. Thin sections were prepared and used for *in situ* hybridization.

Cichlids were anesthetized with ice water for 1 min. The catheter was inserted via the urogenital papilla into the urinary bladder. The silicon tubing connecting the catheter was fixed to the anal fin by a wire and then clipped to hold it in place. Catheterized cichlids were placed in a polyethylene net chamber to restrict the movement. Urine was aspirated through the catheter for 3–5h. Approximately 500–1000 µL of urine was trapped into the centrifuge tube placed on ice. Urine collected during the first 30 min was discarded to prevent contamination by coelomic fluid. To ensure that the sample collected was urine, 10 µL of the collected sample was used to verify the existence of ammonia by indophenol assay (Tetra Test Ammonia Reagent, Tetra). Urine was diluted 30-fold with ultrapure water prior to exposure treatment, and 15 mL diluted urine was applied experimentally (final concentration at 6,000-fold dilution in the exposure tank).

All experimental studies using animals were approved by the Institutional Animal Experiment Committee of the Tokyo Institute of Technology and conducted according to institutional and governmental ARRIVE guidelines.

### Exposure and tissue preparation

Adult cichlids were isolated in a glass tank (40 cm × 25 cm × 36 cm) the day prior to exposure and was not fed. The following day, fish was transferred to the exposure tank (30 cm × 11 cm × 9 cm, 3 L) which was covered with black paper to create a dark environment. Clean dechlorinated water flowed into the tank on one end and out from the opposite end. The fish were kept in this tank for 1.5–3 h before exposure in order to minimize the *c-fos* expression in the OE. Immediately prior to exposure, water inflow was temporarily stopped, and 15 mL of odorant solution was delivered to the same end of the tank as the water inflow using a peristaltic pump (SJ-1211II-H, Atto). Water-only was applied as a negative control. The odorant solution was delivered into the tank over a period of approximately 1 min, and the water inflow was then resumed. Fish were kept in the tank for 20 min after exposure to allow for expression of *c-fos*. The fish were then quickly decapitated and the olfactory epithelia were dissected out in 4% paraformaldehyde (PFA, Wako)/ phosphate-buffered saline (PBS). Dissected tissues were fixed in 4% PFA/ PBS at 4°C for 7.5 h. Fixed tissues were treated with 20% sucrose/ PBS at 4°C overnight for cryoprotection before embedding. Tissues were embedded in the Tissue Tek O.C.T. compound (Sakura) and frozen using liquid nitrogen. Embedded tissues were sliced into a 10 µm horizontal sections and placed on a glass slide (MAS-01, Matsunami). Sections were kept at −80°C until use for downstream experiments. All individuals used in this experiment were male.

### Preparation of riboprobes

Riboprobes for *in situ* hybridization (ISH) were designed in the coding region or untranslated region. Each sequence was amplified from cDNA of the OE by Ex-Taq (Takara) with primers (Supplementary Table 1). PCR products were ligated to pGEM-T (Promega) or pBluescript SKII (−) plasmid and sequenced. Plasmids were extracted with the QIAfilter Plasmid Midi Kit (QIAGEN) and then linearized using an appropriate restriction enzyme (Takara). Digoxigenin (DIG)-labeled or fluorescein (FITC)-labeled riboprobes were synthesized with T7 or T3 or SP6 RNA polymerase (Roche) from the linearized plasmids with DIG or FITC RNA labeling mix (Roche), respectively.

### *In situ* hybridization

Single-colour and two-colour ISH were performed according to the method of Suzuki *et al*., (2015) with several modifications. Briefly, in single-colour ISH, sections were treated with 5 µg/mL proteinase K for 8 min at 37°C and hybridized with DIG-labeled riboprobes (5 ng/µg) at 60°C overnight. The sections were washed, treated with 2 µg/mL RNase A in TNE (Tris-NaCl-EDTA) for 30 min at 37°C, treated with streptavidin/biotin blocking kit (Vector Laboratories), and treated with 1% blocking reagent (PerkinElmer) in TBS (Tris-buffered saline) for 1h. Signals were detected with peroxidase-conjugated anti-DIG antibody (1:100, Roche), amplified by Tyramide Signal Amplification (TSA) Plus Biotin kit (PerkinElmer) and visualized with Alexa Fluor 488-conjugated streptavidin (1:200, Thermo Fisher Scientific). Sections were mounted with VECTASHIELD mounting medium with DAPI (Vector Laboratories). In the case of two-colour ISH, sections were hybridized with DIG-labeled riboprobes and FITC-labeled riboprobe (2.5 ng/µg at each) at 60°C overnight and treated with peroxidase-conjugated anti-DIG antibody (1:100). Signals from DIG-riboprobes were detected with peroxidase-conjugated anti-DIG antibody (1:100, Roche), amplified using TSA Plus DIG kit (PerkinElmer) and visualized with DyLight 594-conjugated anti-DIG antibody (1:500, Vector Laboratories). Sections were treated with 15% H_2_O_2_ in TBS for 30 min to inactivate peroxidase activity before the detection of signals of FITC-labeled riboprobes. Signals from FITC-labeled riboprobes was detected with an anti-Fluorescein-POD, Fab fragments (1:80, Roche), amplified using TSA Plus Biotin kit (PerkinElmer) and visualized with Alexa Fluor 488-conjugated streptavidin (1:200, Thermo Fisher Scientific). Sections were mounted with VECTASHIELD mounting medium with DAPI (Vector Laboratories). All images were digitally captured using the Zeiss Axioplan SP fluorescence microscope with the Zeiss Axiocam 503 color CCD camera. The images were level corrected, contrast adjusted, pseudo-coloured, and merged using Adobe Photoshop CC 2018.

### Analysis of *in situ* hybridization images

For determination of the marker gene of the neural activity, food extract was exposed to cichlids and single ISH (30 min for the time between odorant exposure and ice anesthesia) was performed to find the early marker gene with the greatest increase of positive neurons and the highest signal intensity. For quantification of neurons, a typical shapes of olfactory sensory neurons (ciliated/microvillous/crypt/kappe/pear-shaped neuron; Hamdani and Døving, 2007; Ahuja et al., 2015; Wakisaka et al., 2017) was used as a reference criteria. In addition, for quantification of *c-fos*^+^ neurons, signals which have a luminance value greater than 85 were considered positive. For single colour *in situ* hybridization, 5 sections per individual and 4 individuals were used (Figure 1). For two-colour *in situ* hybridization, one section per individual and four individuals (Figure 3), three individual (Figure 5B, C) or one individual (Figure 4 and Supplementary Figure 1), or 3-7 sections per individual and one individual (data in Figure 5E, G) were used. The area used to correct the number of neurons was measured by the DAPI image of the OE section. For the determination of double-positivity, when both signals were positive, overlapped, and both could be determined to have the same cell shape, we scored the to be double-positive. These procedures were all performed using Adobe Photoshop CC 2018.

### Phylogenetic analysis

All amino acid sequences of *V2R* (East African cichlid [*Haplochromis chilotes*], zebrafish [*Danio rerio*], Atlantic salmon [*Salmo salar*], three-spined stickleback [*Gasterosteus aculeatus*], fugu [*Takifugu rubripes*], green-spotted pufferfish [*Tetraodon nigroviridis*], and medaka [*Oryzias latipes*]) were obtained from Nikaido *et al*. (2013). Sequences were aligned by MAFFT 7 (Katoh and Standley, 2013). RAxML-NG (Kozlov *et al*., 2019) was used to construct an ML tree under the PROTGTR+G+I model with 100 bootstrap replicates.

## Results

### Cichlid *c-fos* has characteristics of an immediate-early gene

We initially exposed cichlids to food extract and assessed the upregulation of five immediate-early genes, *c-fos, egr1, c-jun, fra1*, and *junb* by *in situ* hybridization in the OE. Results of these initial experiments were used to decide on the most suitable gene for neural activity marker detection. Consequuently, *c-fos* was the gene with the greatest increase in signal intensity and number of positive neurons, so *c-fos* was selected as the most suitable neural activity marker for cichlid OE (Figure 2A). To quantitatively confirm the upregulation of *c-fos* in the cichlid OE, we exposed cichlids to food extract and tested whether the number of *c-fos*^+^ neurons would increase with time after exposure (from exposure until ice anesthetizing; control/10/20/30 min) after exposure (Figure 2B, C, Supplementary Table 2). We compared the number of *c-fos*^+^ neurons in five evenly selected sections from each individual and showed that the numbers at 20 min and 30 min intervals was significantly greater in the control (3 individuals; *p* = 0.014, *p* = 0.020; Tukey–Kramer test, Figure 2B, Supplementary Table 3). Thus, we set the time length between odorant exposure and ice anesthetizing as 20 min.

**Figure 2.**
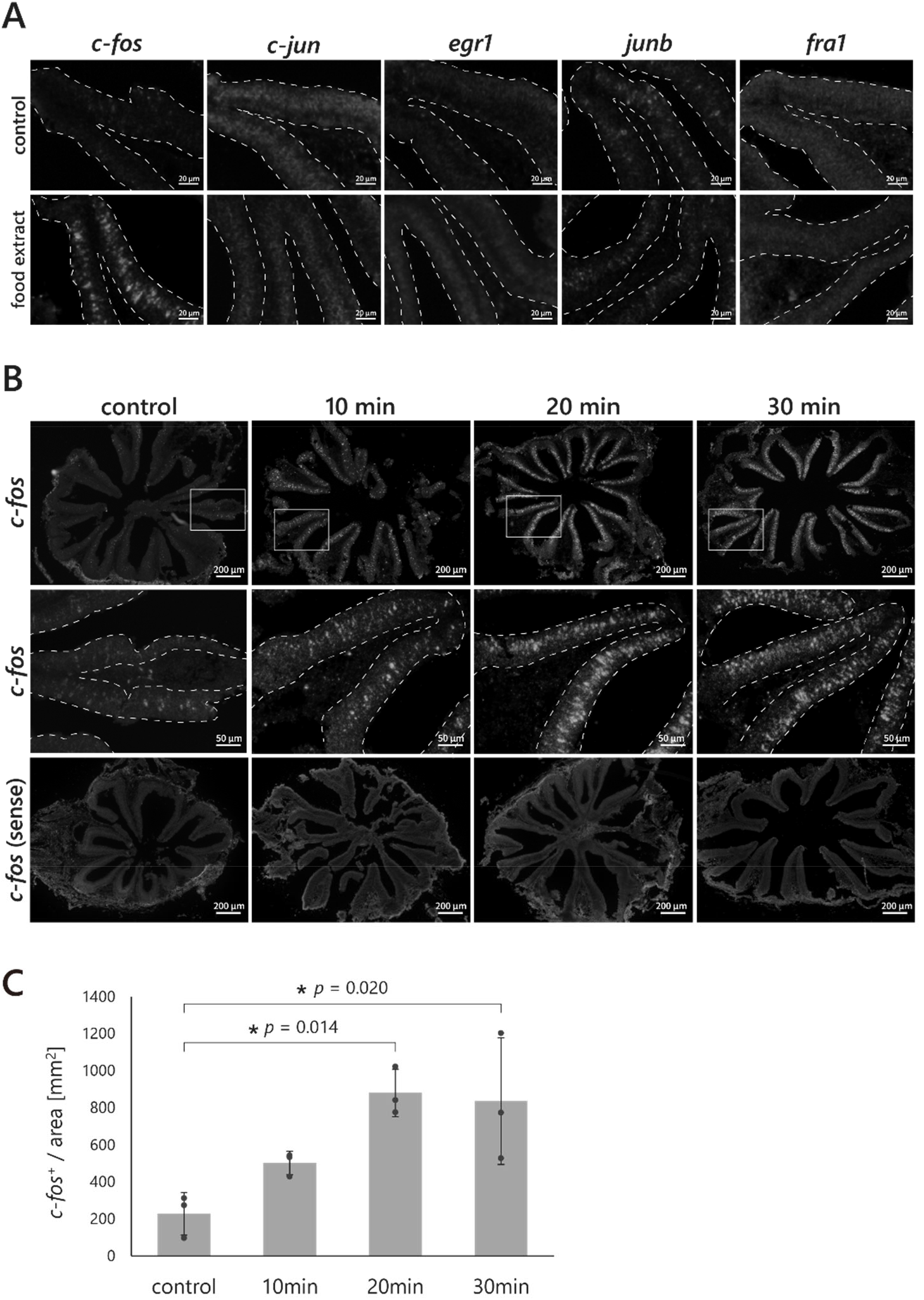
*c-fos* expression is induced robustly sin olfactory epithelium (OE) by exposure to food extract. Cichlids were exposed to water (control) or food extract (final concentration at 15,000-fold dilution). **(A)** *In situ* hybridization with riboprobes for *c-fos, c-jun, egr1, junb*, or *fra1*. The dotted line represents the outline of the OE. **(B)** *In situ* hybridization with riboprobes for *c-fos*. The vertical columns represent the length of time between odorant exposure and ice anesthesia. The middle panels represent the magnified image of the box in the upper panel. **(C)** Bar graph of the number of *c-fos*^+^ in 1 mm^2^. The total number of *c-fos*^+^ neurons in five sections are counted in each individual (three individuals each; Tukey–Kramer test, Supplementary Table 3). \**p* < 0.05. All data are shown as mean ± SEM.

### Neural response of microvillous neurons

Next, we tested the neural response of microvillous neurons. In several fishes, microvillous neurons are known to detect amino acids (Sato and Suzuki, 2001; Hansen et al., 2003; Sato and Sorensen, 2018), and V2Rs, which are expected to be expressed in majority of a microvillous neurons (Supplementary Figure 2A). V2Rs are also suggested to detect amino acids (Koide et al., 2009; DeMaria et al., 2013). In teleosts, amino acids are considered to be associated with food odor. Here, we tested the neural responses of microvillous neurons to the mixture of 20 proteinogenic amino acids (final concentration at 2 µM each) and food extract to examine if our method is useful for testing the specificity of olfactory sensory neurons. In addition, we also tested the neural responses to cichlid male urine (final concentration at 6000-fold dilution) and the mixture of three conjugated steroids (DHEA-s, E2-17g, and E2-3,17s, final concentration at 33 nM each). Urine is considered to be the main source of pheromones in cichlids (Maruska and Fernald, 2012; Keller-Costa et al., 2014). Conjugated steroids are the candidates for cichlid pheromone (Miranda *et al*., 2005; Keller-Costa *et al*., 2014). The three conjugated steroids we tested are known to be detected by independent receptors in African cichlids (Cole and Stacey, 2006). Exposure to the four odorants significantly increased the number of *c-fos*^+^ neurons with strong intensity compared with the control (4 individuals; *p* = 0.047, *p* = 0.035, *p* = 0.024, *p* = 0.043; Welch’s *t*-test, Figure 3A, B, Supplementary Table 2, Supplementary Table 3). We calculated the percentage of *Trpc2*^+^ neurons, which indicate microvillous neurons (Sato *et al*., 2005) among *c-fos*^+^ neurons to test the contribution of microvillous neurons to the detection of each odorant (4 individuals each). The percentage of *Trpc2*^+^ neurons among *c-fos*^+^ neurons became the highest when exposed to amino acids (55% ± 8.8) and was significantly higher than when exposed to male urine (35% ± 5.7) or conjugated steroids (24% ± 3.0) (4 individuals; *p* = 0.0080, *p* = 6.1 × 10^−4^; Student’s *t*-test; Figure 3C, Supplementary Table 2, 3). It also became high when exposed to food extract (48% ± 11; Figure 3C, Supplementary Table 2). Alternatively, it became the lowest when exposed to conjugated steroids and was significantly lower than when exposed to amino acids, food extract, or male urine (*p* = 6.1 × 10^−4^, *p* = 0.0095, *p* = 0.022; Figure 3C, Supplementary Table 3). We further tested the neural responses (one individual each) of another major type of OSNs, ciliated neurons, which can be indicated by *Golf2* expression (Jones and Reed, 1989; Koide et al., 2009). In contrast to microvillous neurons, the percentage of *Golf2*^+^ neurons among *c-fos*^+^ neurons was the highest (67%) when exposed to conjugated steroids, and it became lower in amino acids (42%) and food extract (41%) vs other treatments (58% in male urine) except in control which was 29% (Supplementary Figure 1A, B, Supplementary Table 2).

**Figure 3.**
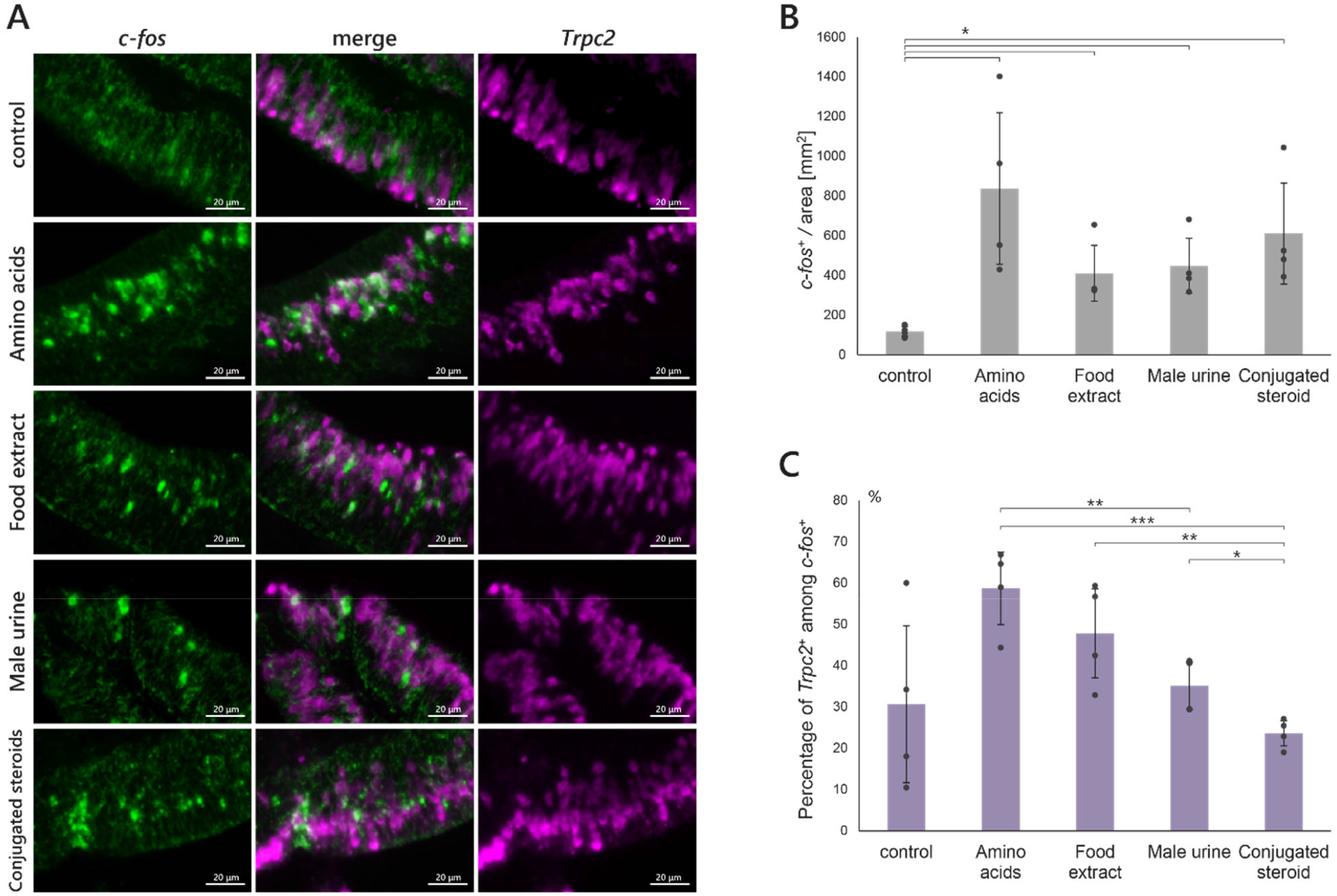
Specificity of microvillous neurons. **(A–C)** Two-colour *in situ* hybridization with riboprobes for *c-fos* (green) and *Trpc2* (magenta) of OE sections of cichlids exposed to water (control), mixture of 20 proteinogenic amino acids (final concentration: 2 µM each), food extract (final concentration: 15,000-fold dilution), male urine (final concentration: 6000-fold dilution), or mixture of three conjugated steroids (final concentration: 33 nM). **(B)** Bar graph of the number of *c-fos*^+^ in 1 mm^2^ (four individuals each; Welch’s *t*-test, Supplementary Table 3). **(C)** Bar graph of the percentage of *Trpc2*^+^ neurons among *c-fos*^+^ (four individuals each; Welch’s *t*-test, Student’s *t*-test, Supplementary Table 3). \**p* < 0.05, \*\**p* < 0.01, \*\*\**p* < 0.001. All data are shown as mean ± SEM.

### Neural response of Vomeronasal type-2 receptor (V2R)^+^ neurons to amino acids

East African cichlids experienced a lineage-specific expansion in the *V2R* multigene family and possess 61 intact *V2R* genes, one of the largest repertoire among teleosts (Nikaido *et al*., 2013). It can be hypothesized that this expanded number of V2R led to the expansion of detectable odors. Cichlids have 13 of the 16 subfamilies of the teleosts previously defined (Hashiguchi and Nishida, 2006), which are composed of 61 *V2R*s (Supplementary Figure 3). Within these 13 subfamilies, especially four subfamilies (4, 8, 14, and 16) have expanded the number of genes by tandem duplication. We therefore tested the neural response of these four expanded subfamilies (4, 8, 14, and 16) plus 2-1, 7-1 as a single-copy subfamily, with a mixture of 20 proteinogenic amino acids (final concentration at 2 µM each, Figure 4A, B, Supplementary Table 2). We designed the riboprobes for each subfamily to have >80% homology with every gene in each subfamily. We made sure in advance that these four subfamilies would not colocalize each other by two-colour *in situ* hybridization (Supplementary Figure 5). The responding rate of *V2R*^+^ neurons was calculated from the percentage of *c-fos*^+^ neurons among *V2R*^+^ neurons. Consistent with the hypothesis that teleosts detect amino acids via V2R receptors, a large fraction of *V2R*^+^ neurons responded to amino acids (Figure 4A, B, Supplementary Table 2). Relatively larger fractions of neurons responded in subfamily 14, 16 (28%/16%), and an intermediate fraction of neurons responded in subfamily 2/7 (8.8%). Alternatively, only a small fraction of neurons responded to amino acids in subfamily 4/8 (3.6%/0.9%).

**Figure 4.**
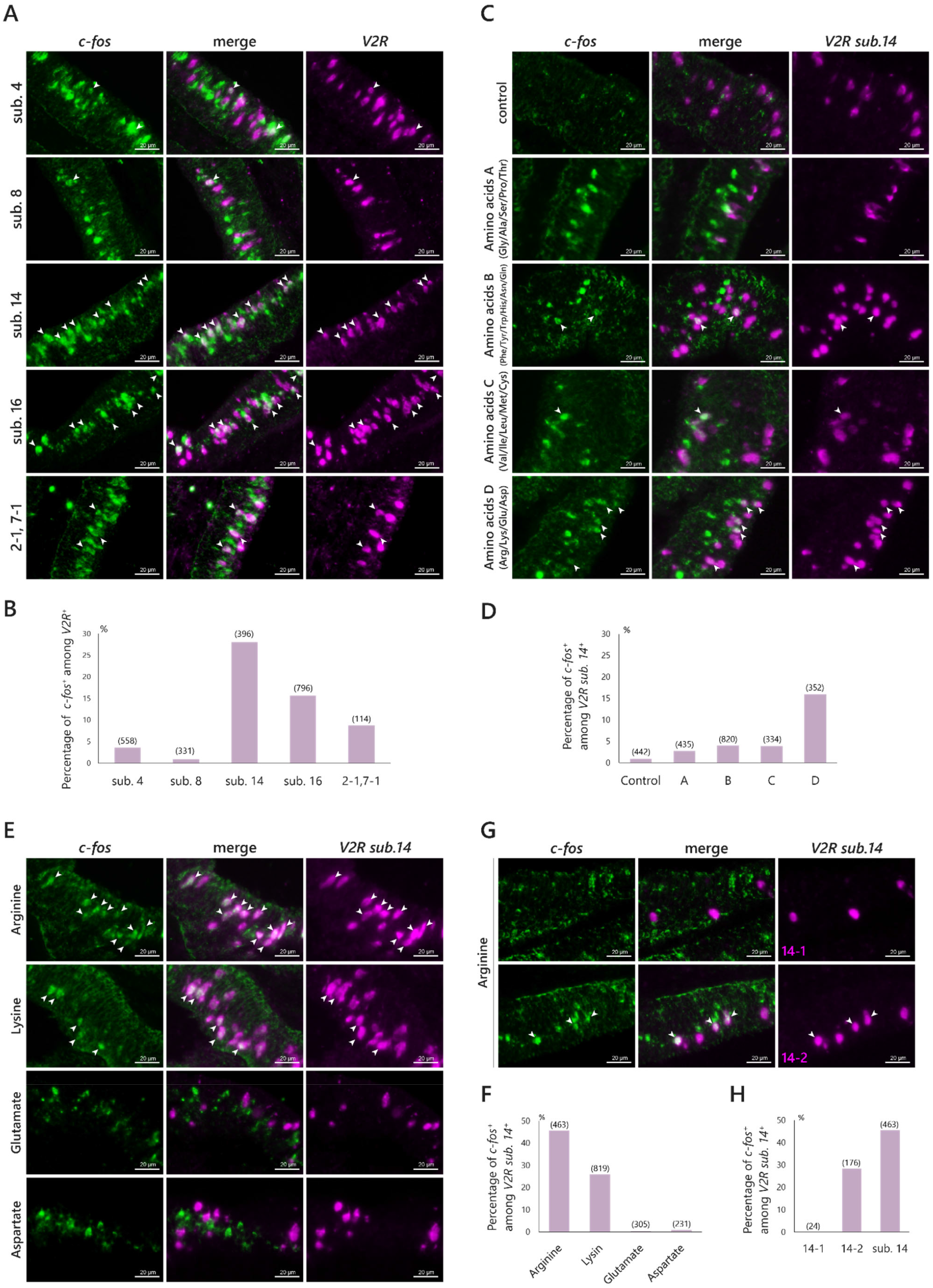
Specificity of *V2R*^+^ neurons to amino acids. **(A, B)** Two-colour *in situ* hybridization with riboprobes for *c-fos* (green) and *V2R subfamily 4/8/14/16/2-1,7-1* (magenta) of OE sections exposed to a mixture of 20 proteinogenic amino acids (final concentration at 2 µM each). **(C, D)** Two-colour *in situ* hybridization with riboprobes for *c-fos* (green) and *V2R subfamily 14* (magenta) of OE section exposed to four groups of amino acids A-D (A: Gly, Ala, Ser, Pro, Thr; B: Phe, Tyr, Trp, His, Asn, Gln; C: Val, Ile, Leu, Met, Cys; D: Arg, Lys, Asp, Glu; final concentration: 2 µM each), arginine (final concentration at 10 µM) or lysine (final concentration at 10 µM). **(E, F)** Two-colour *in situ* hybridization with riboprobes for *c-fos* (green) and *V2R subfamily 14* (magenta) of OE section exposed to arginine, lysine, glutamate, or aspartate (final concentration at 2 µM). **(G, H)** Two-colour *in situ* hybridization with riboprobes for *c-fos* (green) and *V2R 14-1/14-2* (magenta) of OE section exposed to arginine (final concentration at 2 µM). **(A, C, E, G)** Representative image. Arrowheads represent colocalization of *c-fos* and *V2R*. **(B, D, F, H)** Bar graph of the percentage of *c-fos*^+^ neurons among *V2R*^+^ neurons (one individual each). Numbers in brackets represent the number of *V2R*^+^ counted in a single section.

To determine which amino acids are detected by V2R subfamilies 14 and 16, we exposed cichlids to four groups of proteinogenic amino acids: A, including nonpolar or neutral amino acids (Gly, Ala, Ser, Pro, and Thr); B, including aromatic or carbamic amino acid (Phe, Tyr, Trp, His, Asn, and Gln); C, including branched or sulfur-containing amino acids (Val, Ile, Leu, Met, and Cys); and D, including charged amino acids (Arg, Lys, Asp, and Glu) (final concentration at 2 µM each). This grouping is based on electrical properties and a cluster analysis of zebrafish odorant-induced activity patterns (Friedrich and Korsching, 1997). The largest fraction (16%) of *V2R subfamily 14*^+^ neurons responded with stronger intensity to amino acids in D, including charged amino acids (Figure 4C, D, Supplementary Table 2). Although *subfamily 14*^+^ neurons also responded to other amino acid groups, the responding rate was much lower (control: 0.9, A: 2.3%, B: 4.0%, C: 3.9%; Figure 4C, D). On the other hand, the largest fraction of *subfamily 16*^+^ neurons responded to amino acids in group C which including branched or sulfur-containing amino acids, and a small fraction responded to other amino acid groups (control: 0.12%, A: 2.9%, B: 2.8%, C: 9.2%, D: 4.7%; Supplementary Figure 4A, B, Supplementary Table 2).

To further narrow down the amino acids which induce a response in *subfamily 14*^+^ neurons, we exposed cichlids to individual four amino acids in group D (final concentration at 10 µM) and found that basic amino acids, especially arginine, produce strong response in *subfamily 14*^+^ neurons (Arginine: 46%, Lysine: 26%, Glutamate: 0.33%, Aspartate: 0.87%; Figure 4E, F). We further tested the response for arginine of two single-copy genes in subfamily 14 (14-1, 14-2) which are expressed in mutually exclusive manner (Supplementary Figure 5). The response rates to arginine between these two genes were substantially different at 0% and 28%, respectively (Figure 4G, H).

We also tested the neural response of *subfamily 16*^+^ neurons to amino acids in group C of three single-copy genes in subfamily 16 (16-1, 16-3, 16-6). However, no colocalization with *c-fos* was observed in any copy (Supplementary Figure 4C, Supplementary Table 2). This suggests that amino acids in group C are detected by OSNs expressing V2Rs other than 16-1/3/6 in subfamily 16.

We also tested the neural response of four subfamilies of *V2R*^+^ neurons (one individual each) to male urine to examine the possibility that V2R subfamilies that do not respond well to amino acids may be involved in social behavior rather than feeding behavior. However, only a small fraction of *V2R*^+^ neurons responded to male urine (subfamily 4: 1.5%, subfamily 8: 1.7%, subfamily 14: 1.6%, subfamily 16: 1.0%; Supplementary Figure 6A, B).

### Neural response of Vomeronasal type-1 receptor (V1R)^+^ neurons

Finally, we tested the response of *V1R*^+^ neurons. Although zebrafish V1R/ORA has been shown to detect 4HPAA and bile acids (Behrens et al., 2014; Cong et al., 2019), its function remains unclear. We first tested the response of *V1R*^+^ neurons to the four odorants: the mixture of proteinogenic amino acids, food extract, male urine, and the mix of three conjugated steroids using a series of probes for 6 *V1R*s (Figure 5A-C, Supplementary Table 2). *V1R*^+^ neurons responded to male urine with the highest response rate among olfactory stimuli tested (16% ± 5.7; Figure 5A, B), which was significantly higher than that of the control (*p* = 0.038; Tukey-Kramer test; Supplementary Table 3). In contrast, only small fractions of *V1R*^+^ neurons responded to amino acids (5.5% ± 1.3), food extract (5.5% ± 2.8), and conjugated steroids (4.3% ± 3.4) with no significance from the control. Moreover, although the percentage of *V1R*^+^ neurons among *c-fos*^+^ neurons when exposed to male urine was not significantly higher than that of the control (*p* = 0.067), it was significantly higher than that of amino acids, food extract, and conjugated steroids (*p* = 0.026, *p* = 0.030, *p* = 0.028; Tukey-Kramer test; Supplementary Table 3).

**Figure 5.**
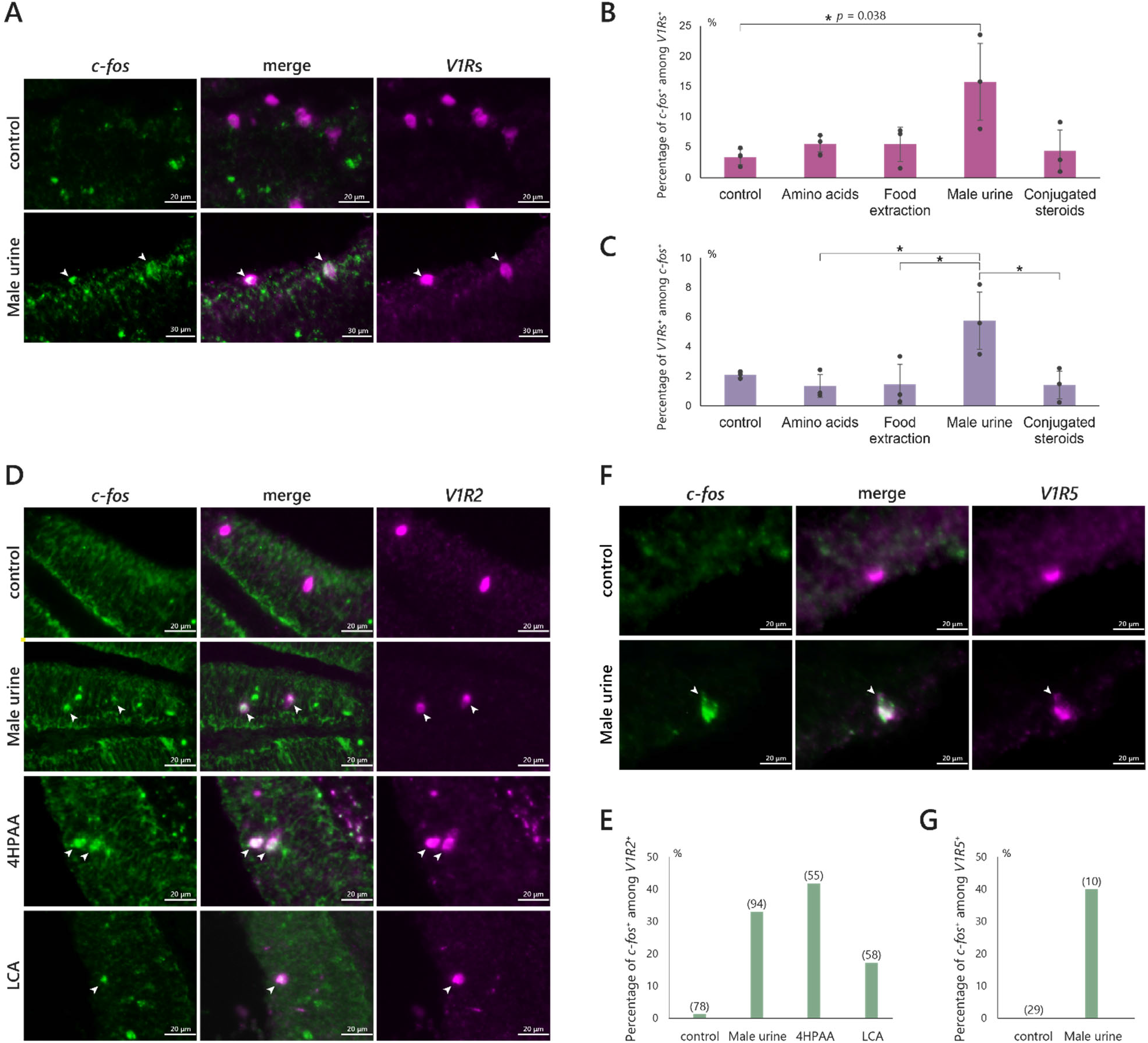
Response of V1R receptor. **(A)** The catheter was used for urine collection. These catheters have a hole in the side near the tip of the needle to prevent clogging. The catheter for females has a silicone plug attached to it. **(B)** Schematic drawing of urine collection. **(C–F)** Double *in situ* hybridization with cRNA probes for *c-fos* (green) and *V1R* (magenta) of olfactory epithelium section. **(C, E, F)** Representative image. Arrowheads represent *c-fos* and *V1R* colocalization. **(D, G)** Bar plot of overlap rate of f *c-fos*^+^ in *V1R*^+^ (1 individual each). The numbers in the brackets represent the number of *V1R*^+^ counted in a single section. **(C, D)** Response of V1Rs (V1R1-6) receptor to amino acids (final concentration: 2 µM each), food extraction (final concentration: 9.5 mg/L), male urine (final concentration: 6000-fold dilution), or conjugated steroids (final concentration: 33 nM). **(E)** Response of V1R2 receptor to male urine (final concentration: 6000-fold dilution), 4HPAA (final concentration: 10 µM), or LCA (final concentration: 10 µM). **(F)** Response of V1R5 receptor to male urine (final concentration: 6000-fold dilution).

We next tested the response to male urine for each of the 6 *V1R*s and found only *V1R2/ORA1*^+^ neurons and *V1R5/ORA5*^+^ neurons were responsive (33%, 40%; Figure 5D-G Supplementary Table 2). Additionally, we tested the response of *V1R2/ORA1*^+^ neurons to 4HPAA (final concentration at 10 µM) and LCA (final concentration at 20 µM), which are the candidates for the ligand of zebrafish V1R2/ORA1 (Behrens *et al*., 2014; Cong *et al*., 2019). Many *V1R2/ORA1*^+^ neurons responded to 4HPAA (42%), and smaller fraction responded to LCA (17%) (Figure 5D, E, Supplementary Table 2).

## Discussion

### Odorant-induced neural responses of cichlid OSN can be tested by *in situ* hybridization with *c-fos* riboprobe

Here, we demonstrated that *in situ* hybridization with riboprobes of *c-fos* are useful for testing the odorant-induced neural responses of cichlid OSNs. Although *egr1*, another major marker gene for neural activity (Isogai *et al*., 2011), was reported to be helpful as an active marker in the cichlid brain (Burmeister and Fernald, 2005), it did not show obvious upregulation in OE. *c-fos*^+^ neurons were significantly increased 20 min after exposure to food extract and the intensity of *c-fos* signals was stronger after 20 min. Furthermore, a large fraction of *Trpc2*^+^ neurons and *V2R*^+^ neurons responded to amino acids, which support previous studies that demonstrate teleosts can detect amino acids via microvillous neurons and V2Rs (Sato and Suzuki, 2001; Hansen et al., 2003; Koide et al., 2009; DeMaria et al., 2013; Sato and Sorensen, 2018). On the other hand, 41% of *c-fos*^+^ neurons were not *Trpc2*^+^ (Figure 3C), and we also implied that 42% of *c-fos*^+^ neurons were *Golf2*^+^ (Supplementary Figure 1B) when exposed to amino acids, which suggests that ciliated neurons also respond to amino acids. This is consistent with previous electrophysical research on rainbow trout, channel catfish, and goldfish (Sato and Suzuki, 2001; Hansen *et al*., 2003; Sato and Sorensen, 2018). These results also suggest that the upregulation of *c-fos* is induced by odorant exposure.

Large fractions of *V2R subfamily 14*^+^ and *subfamily 16*^+^ neurons responded to amino acids, which supports previous studies (Koide et al., 2009; DeMaria et al., 2013). Alternatively, we showed that *V2R subfamilies 4*^+^ and *8*^+^ neurons only marginally responded to proteinogenic amino acids. This indicates that a majority of V2R receptors in subfamilies 4 and 8 receive other chemical compounds, such as nonproteinogenic amino acids and peptides. Kynurenine is one example of a nonproteinogenic amino acid that Masu salmon detect as a sex pheromone (Yambe et al., 2006). Mouse V2R receptors recognize peptides (Kimoto *et al*., 2005), and stickleback and zebrafish detect 9-mer MHC peptides via the OE (Milinski *et al*., 2005; Hinz *et al*., 2013). These chemical compounds are possibly playing a role other than foraging. V2R subfamily 9 has been implicated in the fright reaction of Ostariophysan fishes (Yang et al., 2019). Furthermore, peptides are more suitable for species-specific odor detection since peptides can be more diverse than single amino acids via combination of several amino acid residues. Within teleost *V2R*s, subfamilies 4 and 16 are independently diversified in several lineages (Nikaido *et al*., 2013) suggesting that these subfamilies could possibly receive species-specific odors. V2R subfamily 4 only marginally responded to proteinogenic amino acids and at least three genes of V2R subfamily 16 did not respond to proteinogenic amino acids (Figure 4B; Supplementary Figure 4C).

Also, the *V2R* gene cluster is adjacent to *neprilysin*, which is encoding membrane-bound neural metallopeptidase. Since *neprilysin* is upregulated during ovulation (Langenau et al., 1999), it has been argued that peptides cleaved by neprilysin and the degraded peptides released from spawned eggs may be received by V2R receptors (Hashiguchi and Nishida, 2006; Johnstone et al., 2009; Nikaido et al., 2013). We also hypothesized that some peptides entering the nasal cavity would be cleaved by neprilysin and immediately received by V2R receptors. In fact, we confirmed that *neprilysin* is expressed in the cichlid OE (Supplementary Figure 7). Most *neprilysin* was expressed in cells in the basal region of the OE.

### Duplicated *V2R* genes may help cichlid to detect new odorant

We also showed that among the two V2Rs in subfamily 14, only one V2R was receptive to arginine (Figure 4G, H). Moreover, we showed that some V2Rs in subfamily 16 responded to amino acids in group C, whereas the three V2Rs in subfamily 16 that we tested did not respond to amino acids in group C (Supplementary Figure 4C). The different ligand selectivity in the expanded cichlid-specific subfamily suggests that the specific expansion of V2R led to an expansion of detectable odors that would allow for diversification of diet. Previous studies also supported this hypothesis from the result that the residues predicted to be related to ligand selectivity (Luu et al., 2004; Alioto and Ngai, 2006) were much diverse in cichlid-specifically expanded subfamilies than those of the other teleosts (Nikaido et al., 2013).

### Detection of urine in cichlid OE

In this study, we reported the neural response of OSNs to urine for the first time. We showed that 35% of *c-fos*^+^ neurons, which were induced by the exposure of male urine, were microvillous neurons (*Trpc2*^+^), 58% were ciliated neurons (*Golf2*^+^) and 0% were crypt neurons (*V1R4/ORA4*^+^). We found that ciliated neurons contributed the most to the detection of urine. Conjugated steroids are one possible compound in urine that is detected by ciliated neurons. Conjugated steroids are pheromone candidates in cichlids (Cole and Stacey, 2006). In goldfish, ciliated neurons detect sex steroids, which are structurally similar compounds (Sato and Sorensen, 2018). We also found that *V1R*^+^ neurons do not detect conjugated steroids (Figure 5B).

15.8% of *V1R*^+^ neurons, which collocate *Trpc2* (Supplementary Figure 5), responded to male urine, suggesting that urine-responding microvillous neurons contain *V1R*^+^ neurons. However, the population of *V1R*^+^ neurons was much smaller than *V2R*^+^ neurons, and since the percentage of *V1R*s^+^ neurons among *c-fos*^+^ neurons is 5.8% when exposed to male urine (Figure 5C), it is possible that some *V2R*^+^ neurons are also responding to male urine. Although four *V2R subfamily*^+^ neurons did not respond to male urine (Supplementary Figure 6), this does not preclude the possibility that other V2Rs responded to male urine.

Among six V1R receptors, *V1R2/ORA1*^+^ and *V1R5/ORA5*^+^ neurons responded to male urine (Figure 5D, E). Although V1R receptors other than V1R2/ORA1 and V1R5/ORA5 did not respond to urine, they might be responsible for other odorants such as female urine and feces. Another possibility is that they are used to find food since 9% of expressing neurons responded to food extract (Figure 5B).

V1R2 has a higher number of positive neurons in the OE compared to other V1Rs, suggesting that it is particularly important for urine detection. We demonstrated that *V1R2/ORA1*^+^ neurons responded to 4HPAA and LCA (Figure 5D), and it has been shown previously that cultured cells expressing zebrafish V1R2/ORA1 responded to 4HPAA and bile acids (Behrens *et al*., 2014; Cong *et al*., 2019). Previous research showed that exposure to 4HPAA induces spawning of zebrafish (Behrens *et al*., 2014), and our preliminary experiments confirmed that 4HPAA exists in cichlid urine. Thus, V1R2/ORA1 might be a pheromone (= 4HPAA) receptor that is common across teleosts. Notably, two distinct types of V1R2/ORA1 alleles (Nikaido *et al*., 2014) occur in East African cichlids, and the individuals used in this study had the ancestral allele. Further investigation of the function of the alternative alleles should help illuminate the potential impact of the V1R2/ORA1 receptors on adaptive radiation in cichlid fishes via assortative mating.

## Conclusion

In summary, we demonstrated that *in situ* hybridization with riboprobes of *c-fos* are useful for testing the odorant-induced neural responses of cichlid OSNs by showing that: (1) the number of *c-fos*^+^ neurons increased with odorant exposure; and (2) microvillous neurons responded to amino acids and food extract, which is consistent with previous research on zebrafish. We also showed that (3) that each V2R subfamilies have different responsiveness to amino acids; and (4) there is a difference in response to arginine between two copies in V2R subfamily 14, suggesting that duplication of *V2R* may have led to expansion of detectable odorants in cichlids. Furthermore, we (5) established a new method to collect urine nonlethally from cichlids, and (6) showed various OSNs, including *V1R*^+^ neurons (especially *V1R2* and *V1R5*), responded to male urine. Taken together, results of our study verify the ligand specificity of OSNs to odorants in cichlids, which we anticipate will continue to be revealed experimentally as fundamentally important to adaptive radiation in this extraordinarily biodiverse group of teleost fishes.

## Funding

This work was supported by JSPS KAKENHI [grant numbers 20H03307 and 20KK0167 to M.N.] and the Sasakawa Scientific Research Grant from The Japan Science Society [grant number 2021-4099 to R.K.].

## Conflict of interest

The authors do not have any conflicts of interest.

## Acknowledgements

We thank Mitsuto Aibara for filed collection and curation of cichlids from Tanzania, Tatsuki Nagasawa for helpful advice with experiments, and the Biomaterials Analysis Division, Tokyo Institute of Technology, for DNA sequencing support.

## Supplementary Data

**Supplementary Figure 1.**
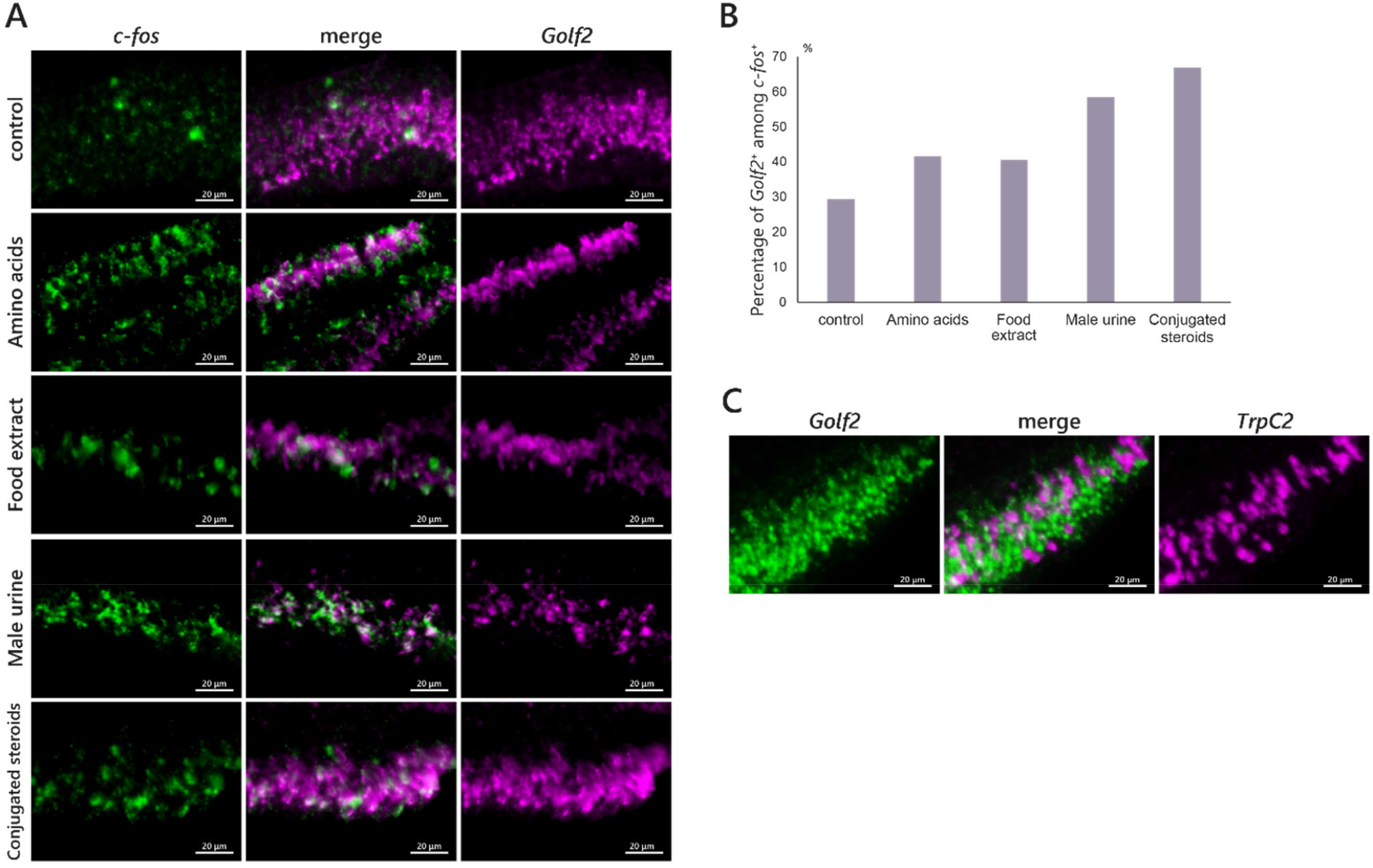
Specificity of ciliated neurons. **(A, B)** Two-colour *in situ* hybridization with riboprobes for *c-fos* (green) and *Golf2* (magenta) of OE sections of cichlids exposed to water (control), mixture of 20 proteinogenic amino acids (final concentration: 2 µM each), food extract (final concentration: 9.5 mg/L), male urine (final concentration: 6000-fold dilution), or mixture of three conjugated steroids (final concentration: 33 nM). **(A)** Representative image. **(B)** Bar graph of the percentage of *Golf2*^+^ neurons among *c-fos*^+^ neurons (one individual each). **(C)** Representative image of two-colour *in situ* hybridization with riboprobes for *Golf2* (green) and *Trpc2* (magenta) of OE sections of cichlids.

**Supplementary Figure 2.**
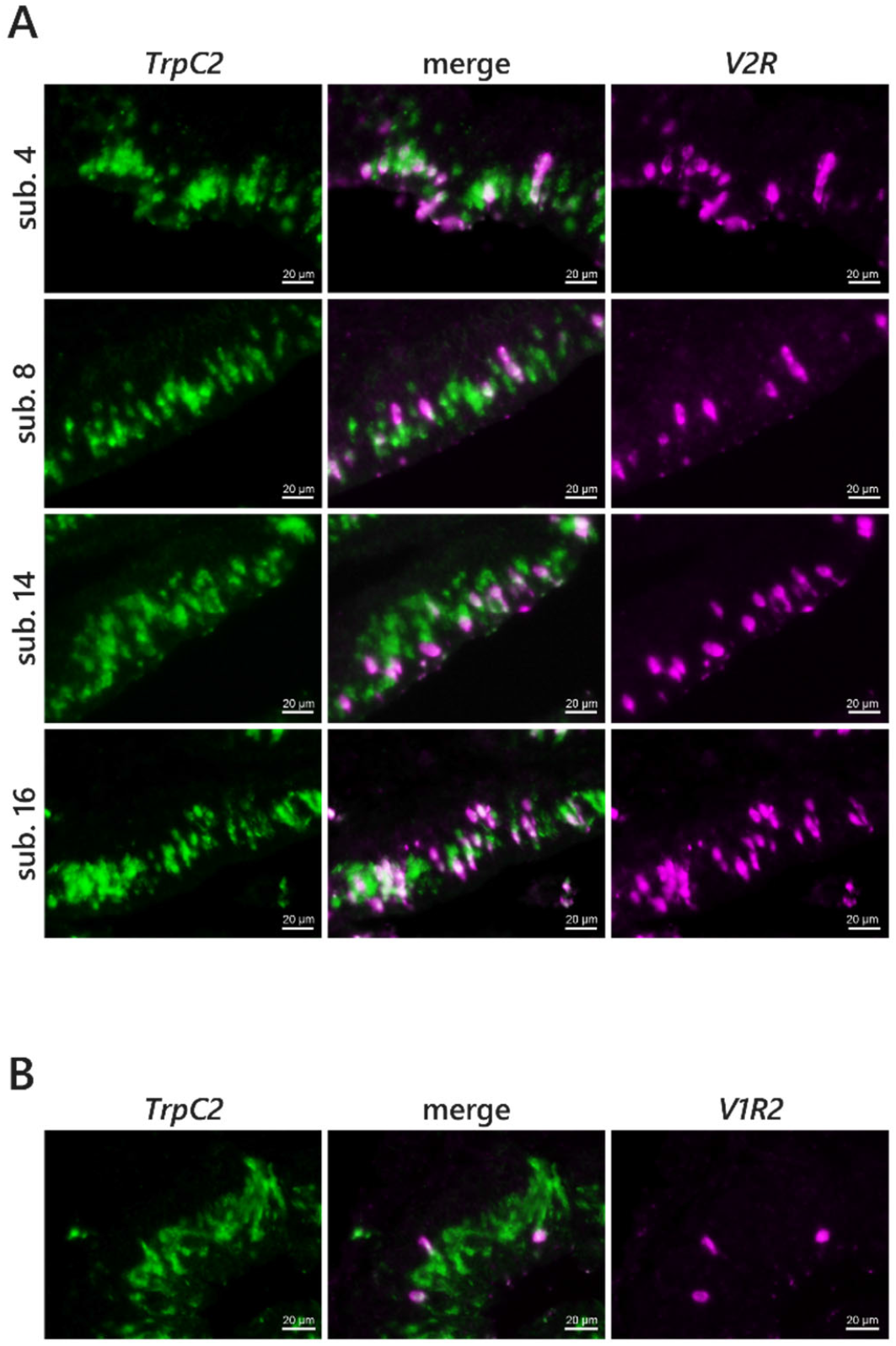
Coexpression of *Trpc2* and *V2R* or *V1R*. **(A-B)** Representative image of two-colour *in situ* hybridization with riboprobes for *Trpc2* (green) and *V2R subfamily 4/8/14/16* (**A**; magenta) or V1R2/ORA1 (**B**; magenta) of OE section. All *V2R*s and *V1R2/ORA1* colocalized with *Trpc2*.

**Supplementary Figure 3.**
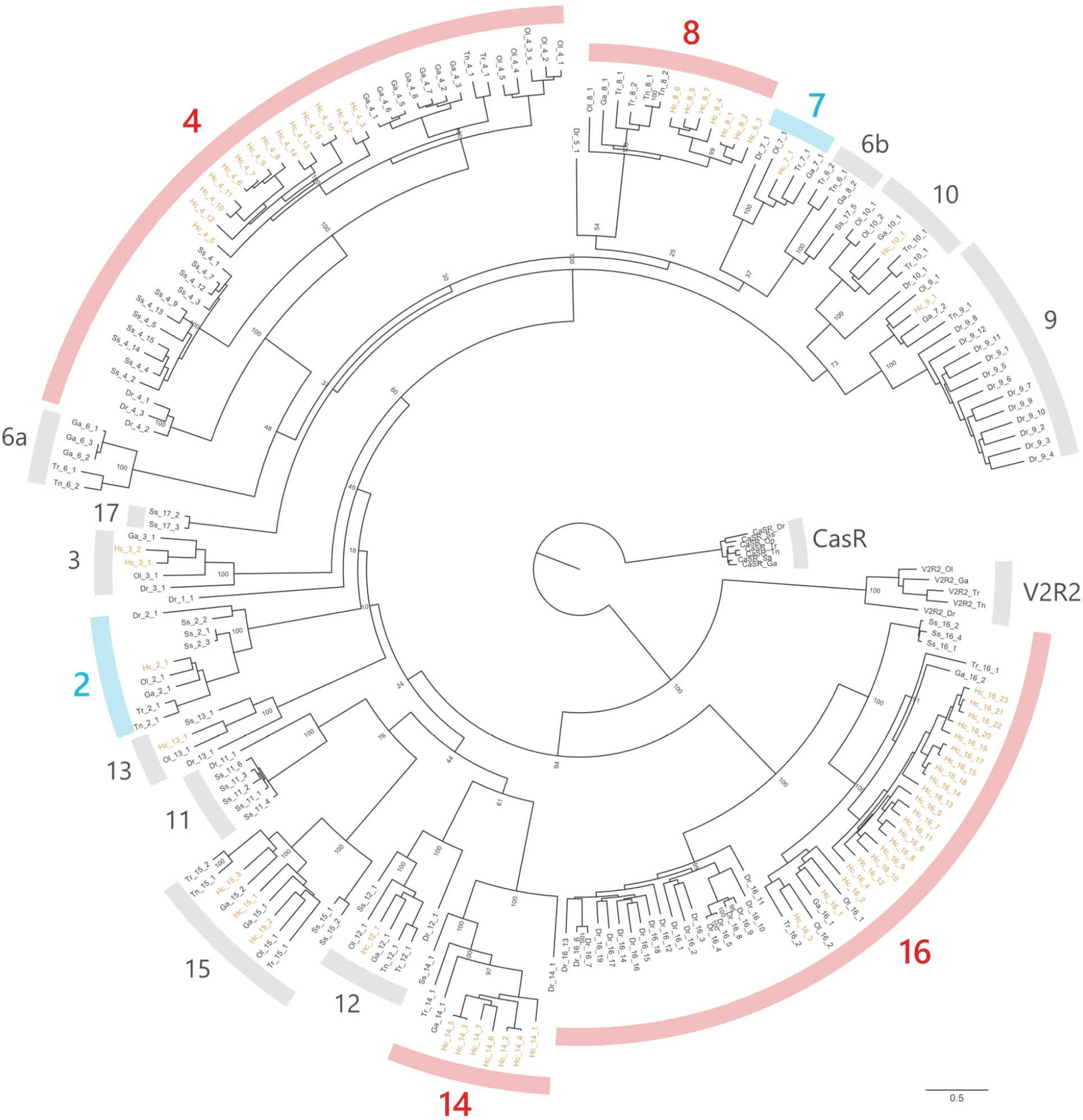
Maximum likelihood (ML) tree for *V2R* genes of seven teleost fishes. All amino acid sequences were obtained from Nikaido *et al*. (2013). RAxML-NG (Kozlov *et al*., 2019) was used to construct an ML tree with 100 bootstrap replicates. Numbers near the nodes are ML bootstrap percentages. Arcs represent each subfamily. Subfamilies used in this study were coloured red (expanded in cichlids) or blue (not expanded in cichlid). Genes of cichlid are coloured orange. Hc: East African cichlid (*Haplochromis chilotes*); Dr: zebrafish (*Danio rerio*); Ss: Atlantic salmon (*Salmo salar*); Ga: three-spined stickleback (*Gasterosteus aculeatus*); Tr: fugu (*Takifugu rubripes*), Tn: green-spotted pufferfish (*Tetraodon nigroviridis*); Ol: medaka (*Oryzias latipes*).

**Supplementary Figure 4.**
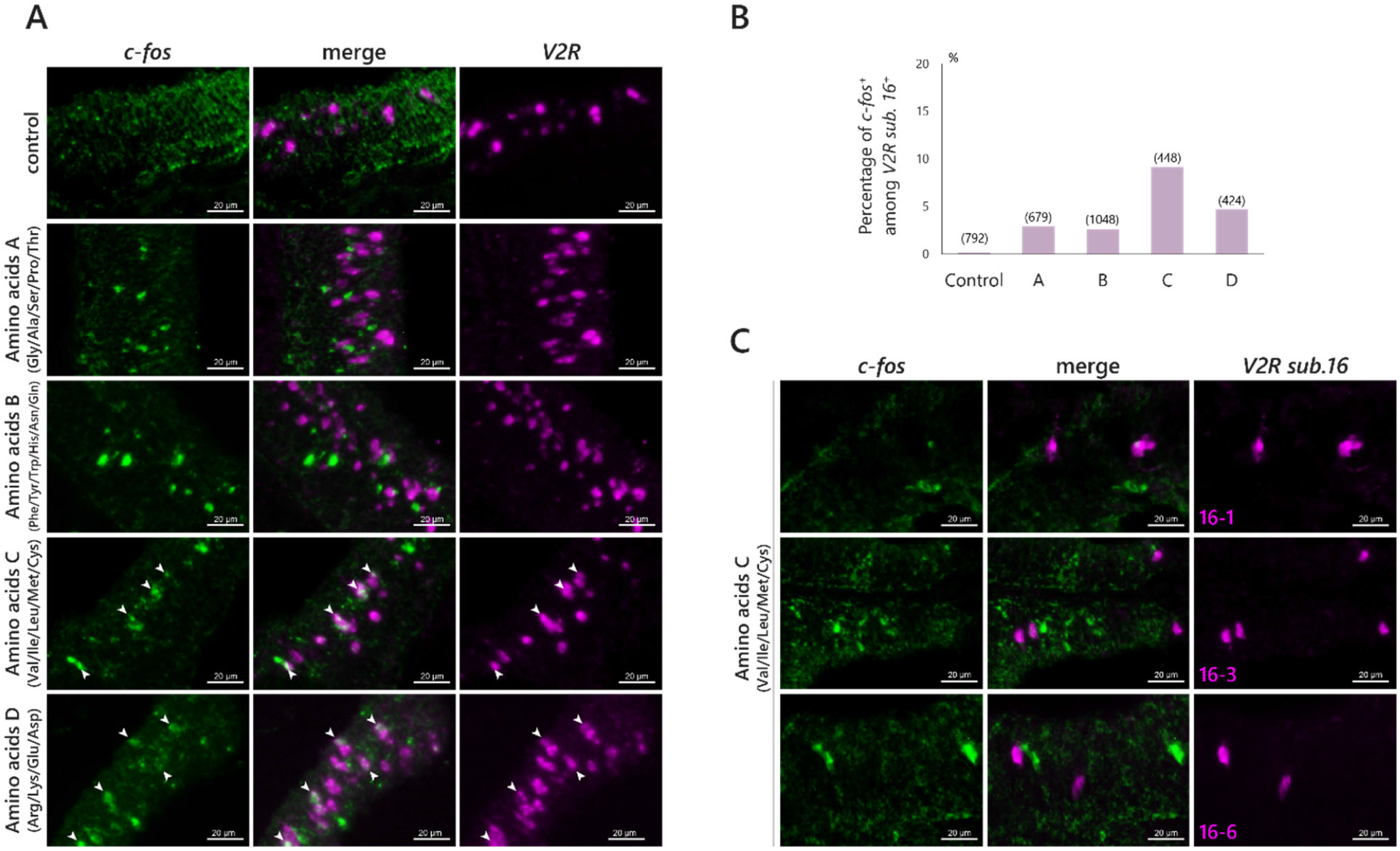
Specificity of *V2R subfamily 16*^+^ neurons to amino acids. **(A, B)** Two-colour *in situ* hybridization with riboprobes for *c-fos* (green) and *V2R subfamily 16* (magenta) of OE sections exposed to four groups of amino acids A-D (A: Gly, Ala, Ser, Pro, Thr; B: Phe, Tyr, Trp, His, Asn, Gln; C: Val, Ile, Leu, Met, Cys; D: Arg, Lys, Asp, Glu; final concentration: 2 µM each), arginine (final concentration at 10 µM) or lysine (final concentration at 10 µM). **(C)** Two-colour *in situ* hybridization with riboprobes for *c-fos* (green) and *V2R 16-1*/*16-3*/*16-3* (magenta) of OE section exposed to amino acids group C (Val, Ile, Leu, Met, Cys; final concentration at 10 µM each) No colocalization was observed. **(A, C)** Representative images. Arrowheads represent colocalization. **(B)** Bar graph of the percentage of *c-fos*^+^ neurons among *V2R*^+^ neurons (one individual each). Numbers in brackets represent the number of *V2R*^+^ counted in a single section.

**Supplementary Figure 5.**
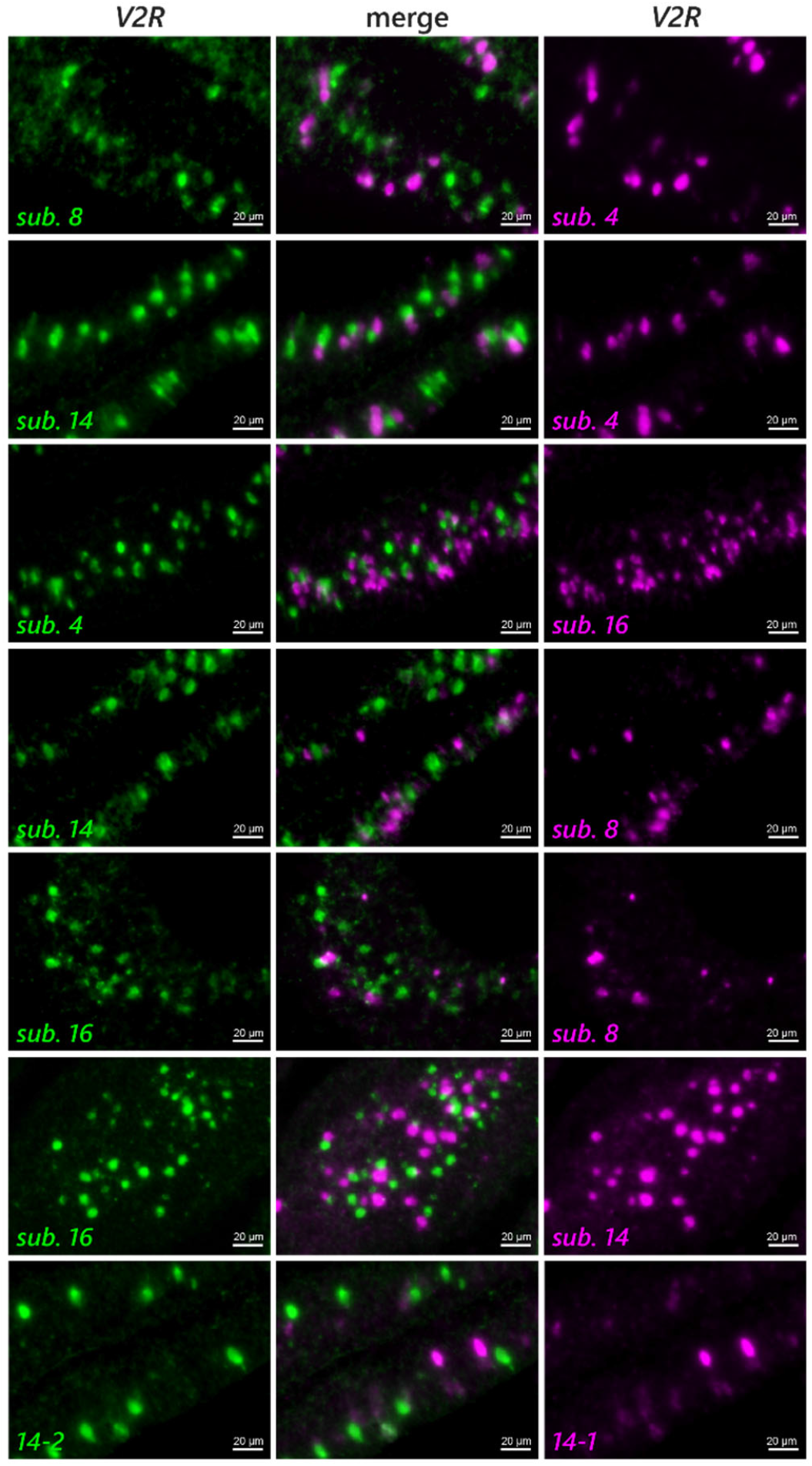
Exclusive expression of *V2R*. Two-colour *in situ* hybridization with riboprobes for two different *V2R*s (green and magenta).

**Supplementary Figure 6.**
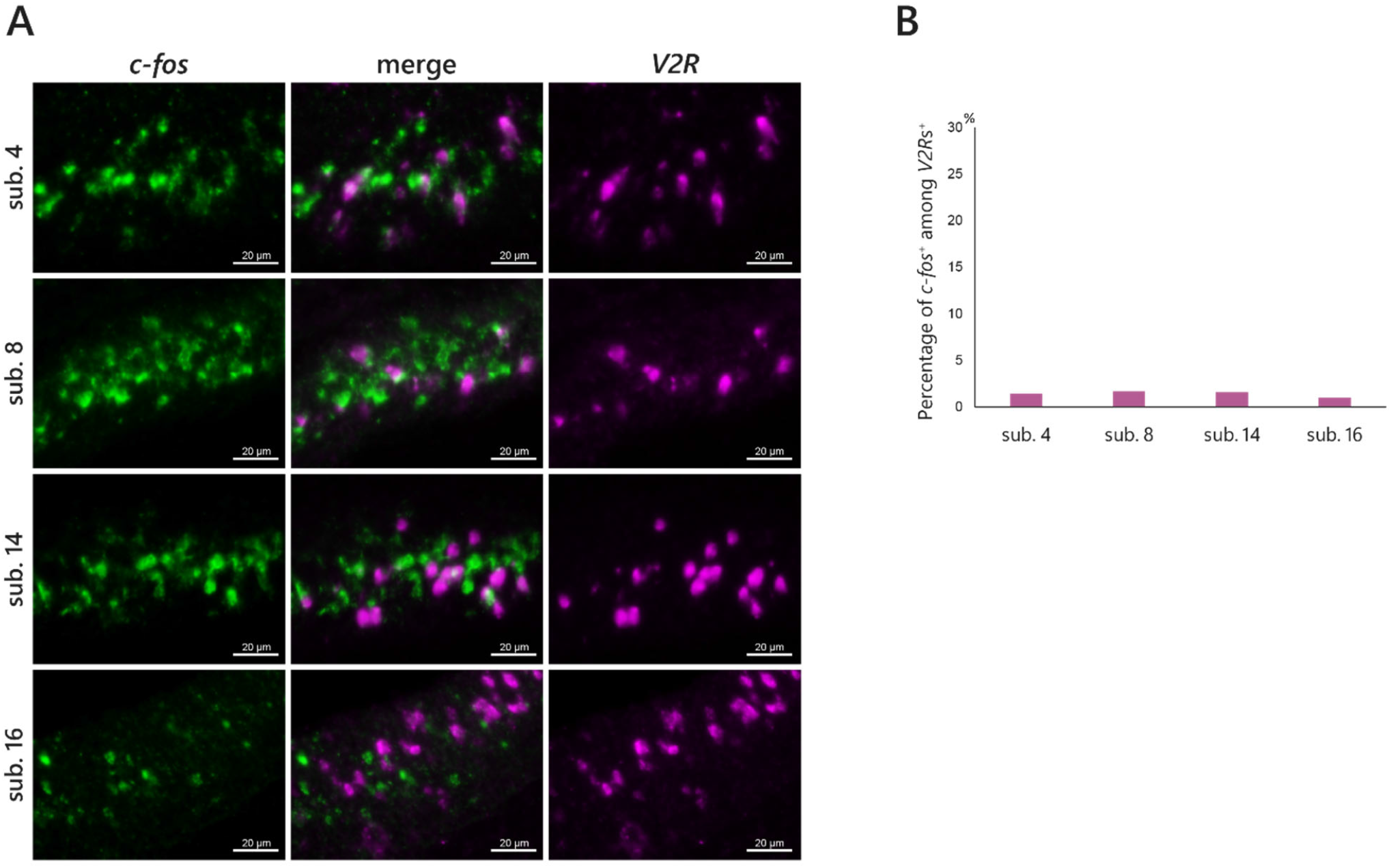
Specificity of *V2R*-expressing neurons to male urine. Two-colour *in situ* hybridization with riboprobes for *c-fos* (green) and *V2Rs* (magenta) of OE sections of cichlids exposed to male urine (final concentration: 6000-fold dilution). **(A)** Representative image. **(B)** Bar graph of the percentage of *c-fos*^+^ neurons among *V2R*^+^ neurons (one individual each).

**Supplementary Figure 7.**
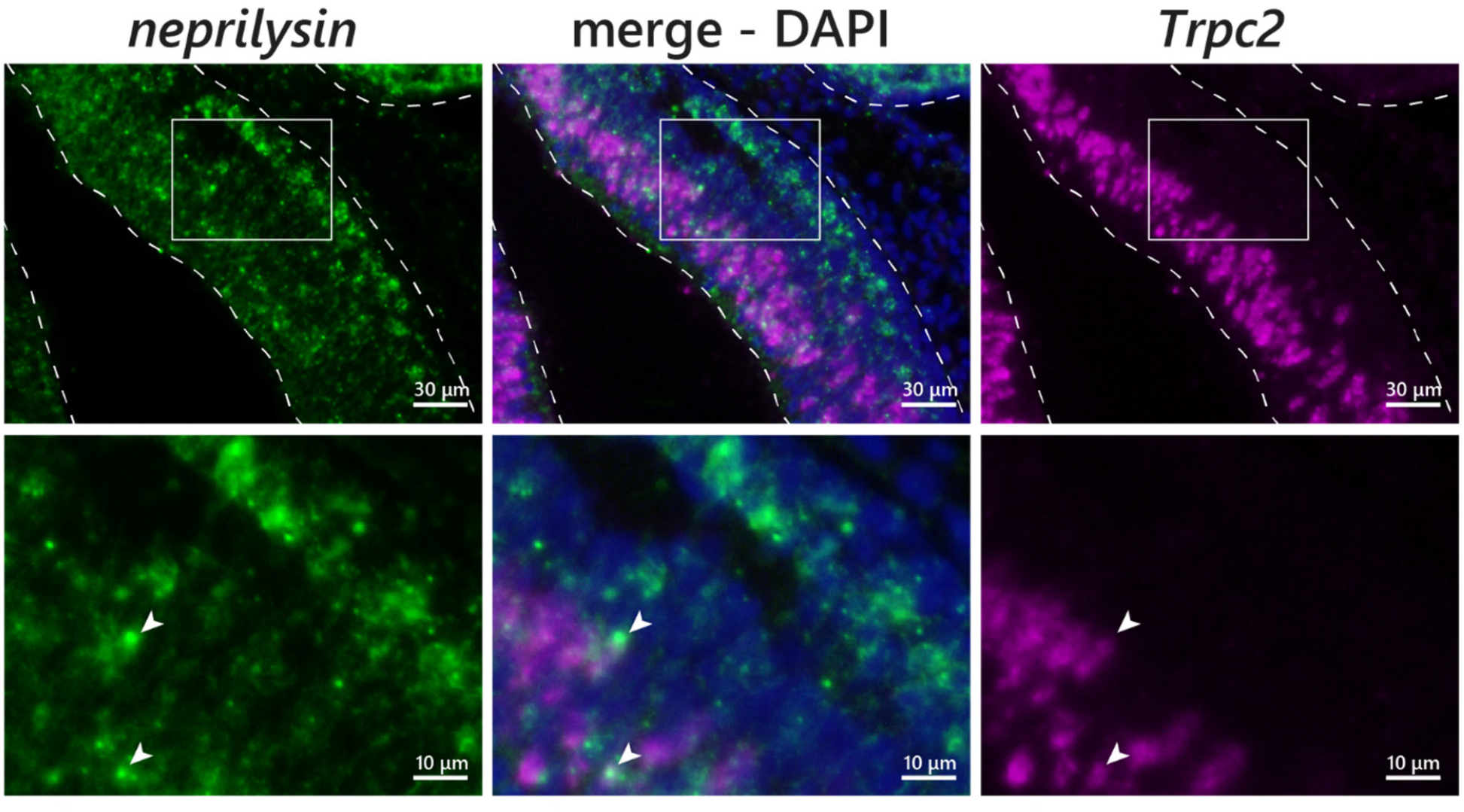
Expression of *neprilysin* in OE. Two-colour *in situ* hybridization with riboprobes for *neprilysin* (green) and *Trpc2* (magenta) of OE sections of cichlids. The middle panels represent the merged image of *neprilysin* and *Trpc2* with DAPI (blue). The dotted line represents the outline of the OE. *Neprilysin* is primarily expressed in the basal region of OE with less expression in the middle region. The lower panels represent the magnified image of the box in the upper panel. Arrowheads represent the colocalization of *neprilysin* and *Trpc2*.

**Supplementary Table 1. Primer list used in this study**.

**Supplementary Table 2. Summary of quantification data analyzed in this study**.

**Supplementary Table 3. Summary of statistical data in this study**.

## Notes

### Competing Interest Statement

The authors have declared no competing interest.

### Summary of Updates

We increased the number of trials on the response of V1Rs neurons to odorants and showed that V1R+ neurons significantly responded to male urine. We also added the results of the response of Golf2+ neurons (ciliated neurons) to odorants (supplementary figure 1) and the response of V2Rs+ neurons to male urine was added (supplementary figure 6). We also revised the manuscript throughout.

